# Bioinformatics evaluation of the homologues of *Schistosoma mansoni* biomarker proteins of bladder cancer in other *Schistosoma* species

**DOI:** 10.1101/2020.09.07.285767

**Authors:** JO Folayowon, AS Adebayo, RD Isokpehi, CI Anumudu

## Abstract

Schistosomiasis remains a public health problem in developing countries. An ideal diagnostic test capable of detecting parasites as early as possible after the onset of infection is therefore highly desired. The identification of biomarker proteins associated with active infection and immune response may constitute the basis for the development of a successful diagnostic test. The aim of this study is to contribute to the development of protein biomarkers for schistosomiasis using a bioinformatics approach. The homologues of 36 previously identified urine biomarker proteins from a *Schistosoma mansoni* database, were identified in other *Schistosoma* species using SMARTBLAST and analyzed for similarities and differences using multiple sequence alignment. Of the 36 *S. mansoni* biomarker proteins, 29 had homologues with both *S. haematobium* and *S. japonicum* or either of *S. haematobium* and *S. japonicum*. Most of the 29 *S. mansoni* biomarker proteins aligned with their homologues with many conserved regions. However, some vital biomarker proteins like venom allergen-like proteins, which had been proposed as a putative drug and vaccine target, showed more semi conserved regions in which the amino acids had similar shape but weakly similar properties. The predictions of 12 markers found in all three species also show that treatment of infections may possibly benefit from the investigational drug Artenimol and specific nutraceuticals or supplements. Experimental evaluation is needed to confirm the potential of the proteins as biomarkers for early diagnosis of schistosomiasis and associated bladder cancer.

## INTRODUCTION

Schistosomiasis, also known as bilharzia, is a disease caused by blood-dwelling trematodes belonging to the genus *Schistosoma*. The genus *Schistosoma* contains six species that are of major pathological importance to man, *Schistosoma haematobium, S. mansoni, S. japonicum, S. mekongi, S. intercalatum*, and *S. guineensis* (French *et al*., 2018). Of these species of flatworms, *S. haematobium S. japonicum* and *S*.*mansoni* are the main schistosome parasites of man (Masamba *et al*., 2016). Schistosomiasis remains one of the most prevalent neglected tropical diseases especially in Nigeria, which supposedly has the greatest number of infected people worldwide (Dawaki *et al*., 2016). Despite the gains in the health care delivery of the past decades, schistosomiasis has prevailed as a health challenge in the tropic and the subtropics. Schistosomiasis is a chronic parasitic infection responsible for nearly 280,000 deaths per year with 200 million people worldwide being infected (Cai *et al*., 2017). According to the World Health Organization (WHO, 2015), schistosomiasis is second to malaria alone amid the vector-borne diseases in terms of public health and remuneration importance in the tropics. An early diagnostic test would enable more rapid treatment and would interrupt the transmission cycle of the parasite and the progress of the disease (Kardoush *et al*., 2016). However, the current diagnostic standards for schistosomiasis all depend on the detection of eggs. In addition, as the disease is asymptomatic in its early stages, clinical examinations cannot confirm infection (Weifang *et al*., 2018).

Human bladder cancer is the fifth to the seventh most common cancer in Western countries. In many tropical and subtropical areas, however, it is the first among all types of cancer, mainly due to the endemic parasitism (Gulia *et al*., 2017). The urinary form of schistosomiasis, known as urogenital or urinary schistosomiasis is caused by *S. haematobium* and is widespread in Africa and the Middle East (WHO, 2015). Epidemiological evidence indicates that *S. haematobium* is associated to squamous cell carcinoma of the bladder. In spite of various control measures and eradication methods that have been in use, schistosomiasis still prevails as one of the most debilitating parasitic diseases, typically affecting the poor and the less privileged predominantly in sub-Saharan Africa (Masamba *et al*., 2016). Efforts to control and eradicate schistosomiasis rely on praziquantel, the only drug available for treatment. However, treatment with praziquantel may sometimes fail, and this may be due to possible drug resistance. There were also case reports of travelers who were infected during their stay in endemic areas and were treated with praziquantel, but resulted in treatment failure (Secor and Montogomery, 2015). Vaccine development against this disease has experienced more failure than successes, (although there is currently a vaccine trial going on for schistosomiasis in Africa (Molehin, 2020); therefore, the identification of molecular targets that can induce protective immunity is highly desirable.

The use of biomarkers in basic and clinical research as well as in clinical practice has become so commonplace that their presence as primary endpoints in clinical trials is now accepted almost without question. The identification of biomarker proteins associated with active infection and protective immune response may constitute the basis for the development of a successful vaccine and could also indicate new diagnostic method (Schmitz-Drager *et al*., 2015). Over the last decade, the number of omics resources used to study this neglected disease has increased considerably, mainly due to advances associated with the development of detection/analysis techniques and mass spectrometry equipment (Hwang *et al*., 2018).

Identification of biomarker proteins is important in early diagnosis of both schistosomiasis and schistosomiasis associated bladder cancer and has been evaluated in recent studies (Onile *et al*., 2017); it may also have application in drug and vaccine development for the treatment of the disease. These biomarker proteins were found using *S. mansoni* database, which is the best curated schistosome species. It is therefore important to know if homologues of these biomarker proteins could be found in other schistosome species, especially in *S. haematobium* and *S. japonicum*. In this present study, we searched for homologues of the thirty-six *Schistosoma mansoni* biomarker proteins found in recent studies in *S. haematobium* and *S. japonicum*. This could be used for early diagnosis of schistosomiasis and schistosomiasis associated bladder cancer. We also attempted to identify proteins common to all *Schistosoma* species which could provide targets for developing drugs or vaccines that can be simultaneously effective against all species of the parasite.

Molecular techniques to detect schistosome infections have been developed to facilitate early diagnosis, but these are expensive and suffer from sampling limitations (Yumin *et al*., 2018; Weerakoon *et al*., 2018). Serologic assays to detect antibodies against schistosome antigens, however, have proven useful in the clinical diagnosis of schistosomiasis. Zheng *et al*., (2012) strongly suggest that SjSP-216, a highly expressed gene in the young worm stage, could serve as a potential biomarker for the early immunodiagnosis of *S. japonicum* infections in vertebrate hosts The aim of the study is to contribute to the development of protein biomarkers for schistosomiasis diagnosis using a bioinformatics approach. This will be achieved by identifying the homologues of the 36 biomarker proteins from *Schistosoma mansoni* evaluated in recent study (Onile *et al*., 2017) in other *Schistosoma* species using bioinformatics tools. These biomarker protein sequences from *S. mansoni* will be compared with those from other *Schistosoma species* to visualize similarities and differences.

## RESULTS

A total of 36 *S. mansoni* biomarker proteins (Table 3.1) were used as query for the homology analysis with *S. haematobium* and *S. japonicum*. Out of these, identical proteins were retrieved for twenty-nine proteins (Table 4.1).

**Table 1:**
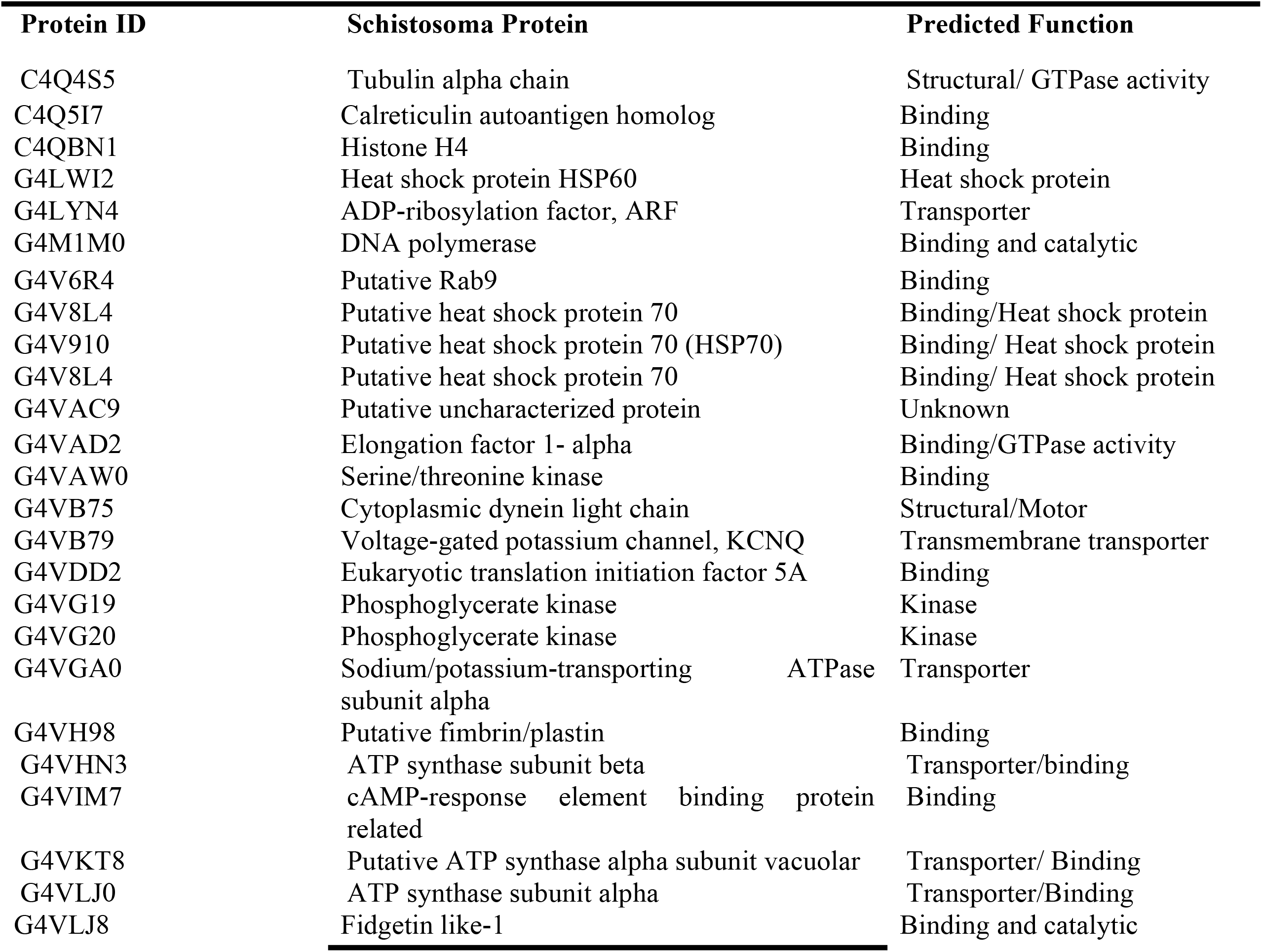
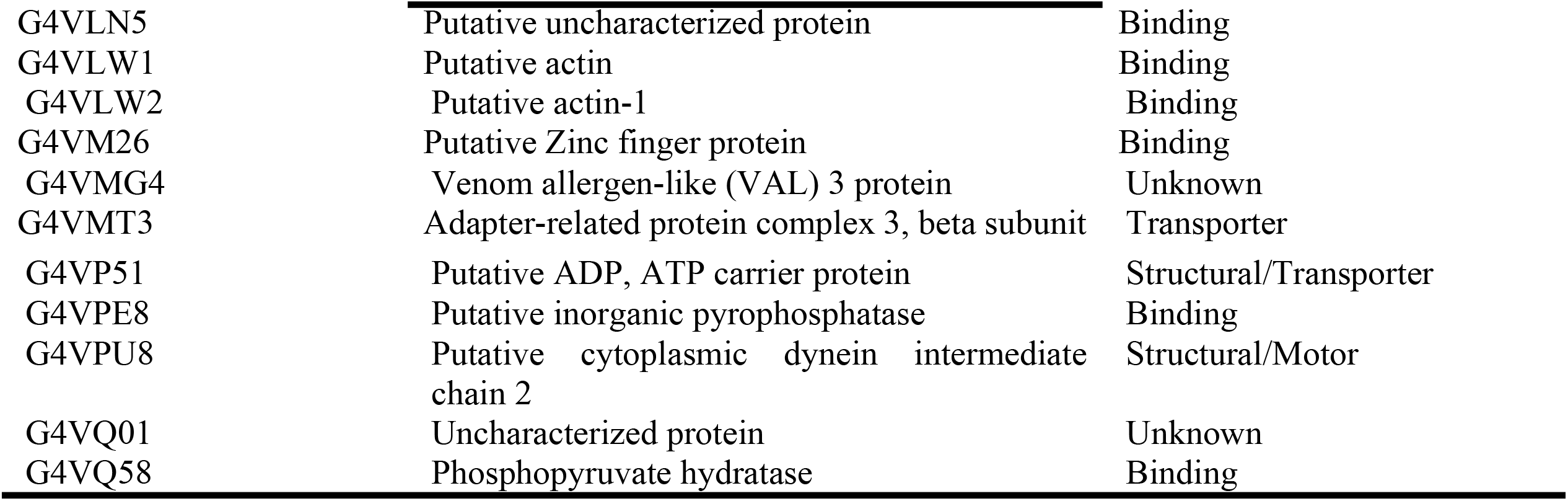
The 36 identified *Schistosoma mansoni* biomarker proteins retrieved from Onile *et al*., 2017 and their predicted function.

**Table 2:**
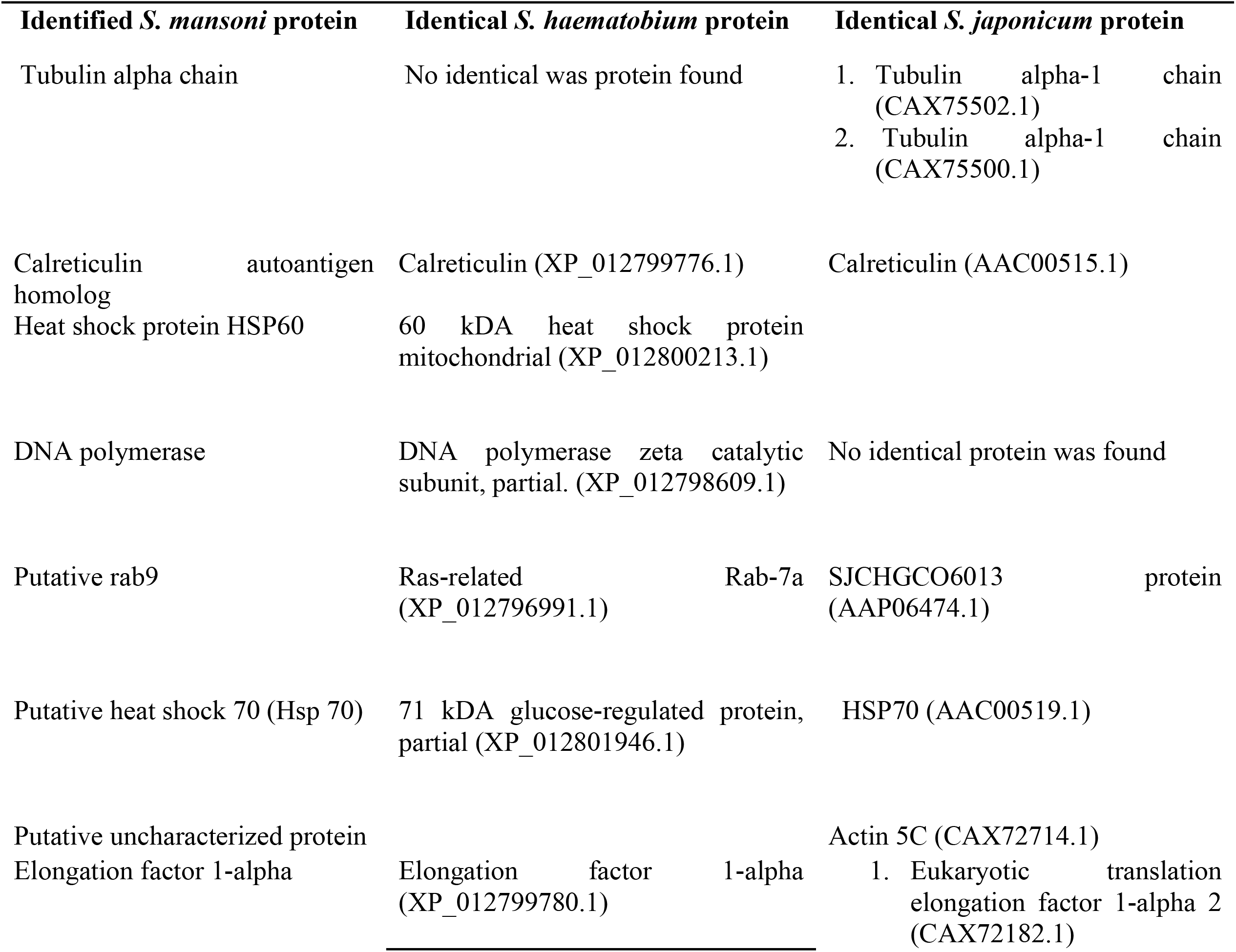
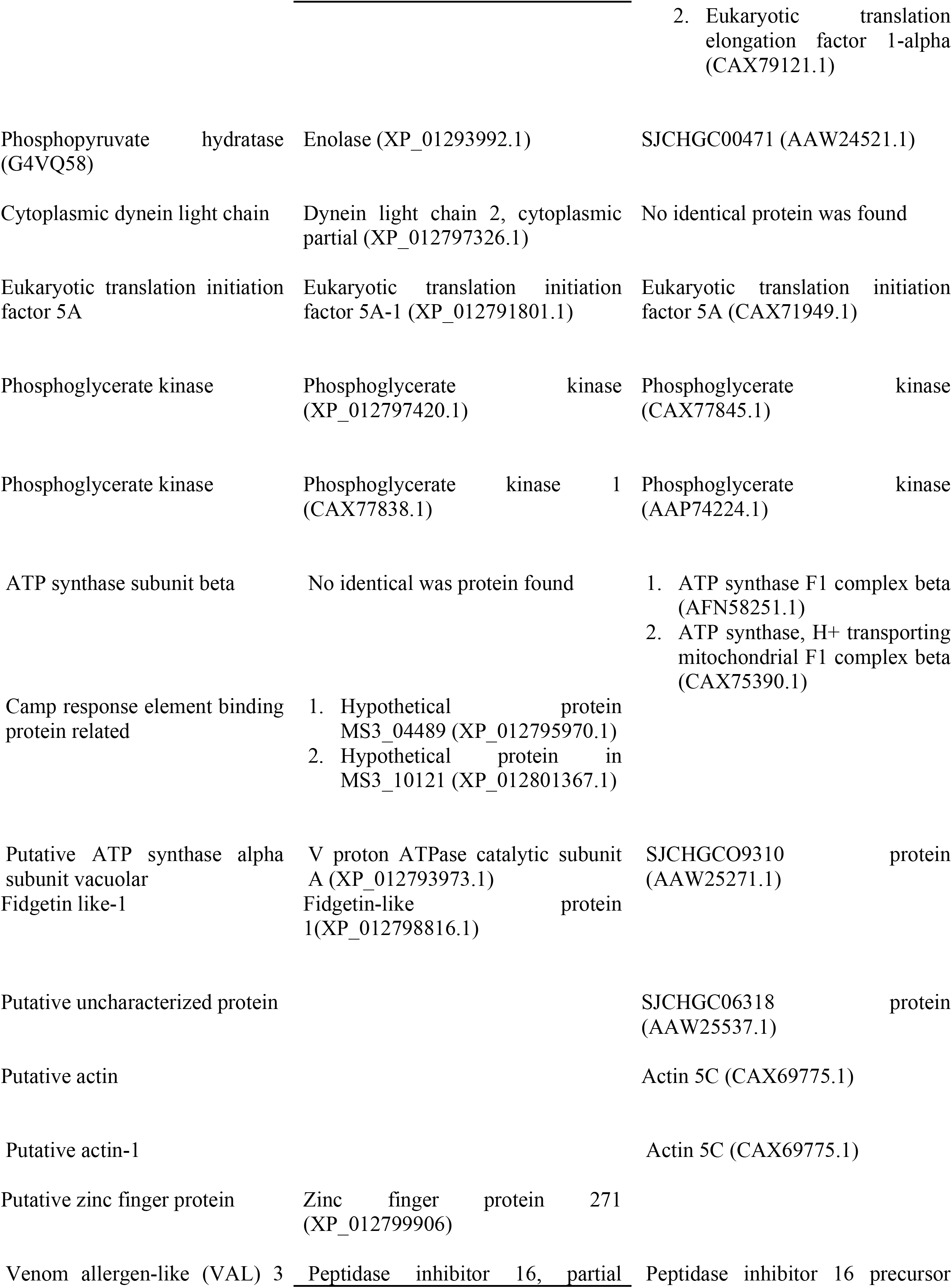
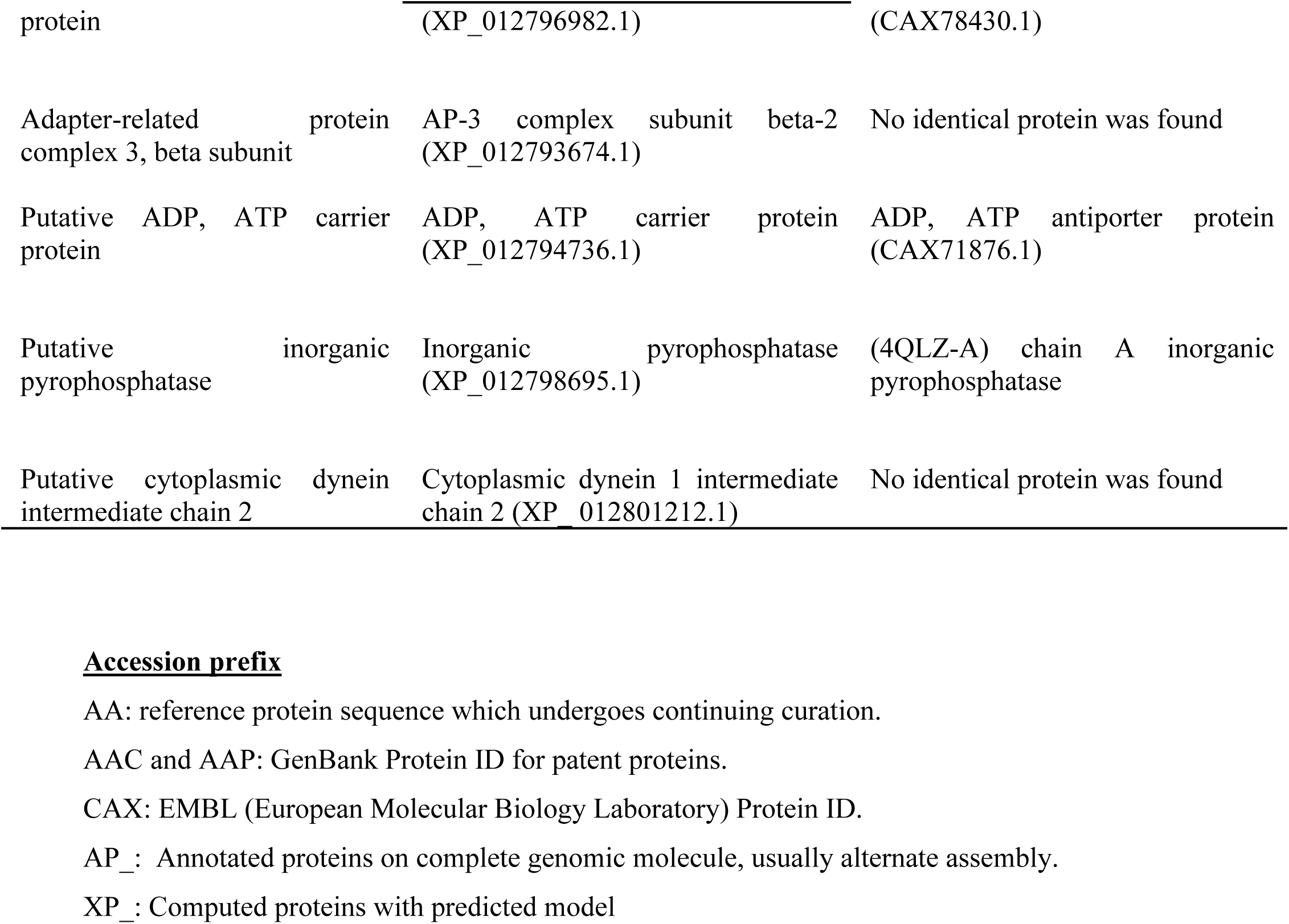
*Schistosoma mansoni* proteins and their retrieved identical proteins

**Table 3.**
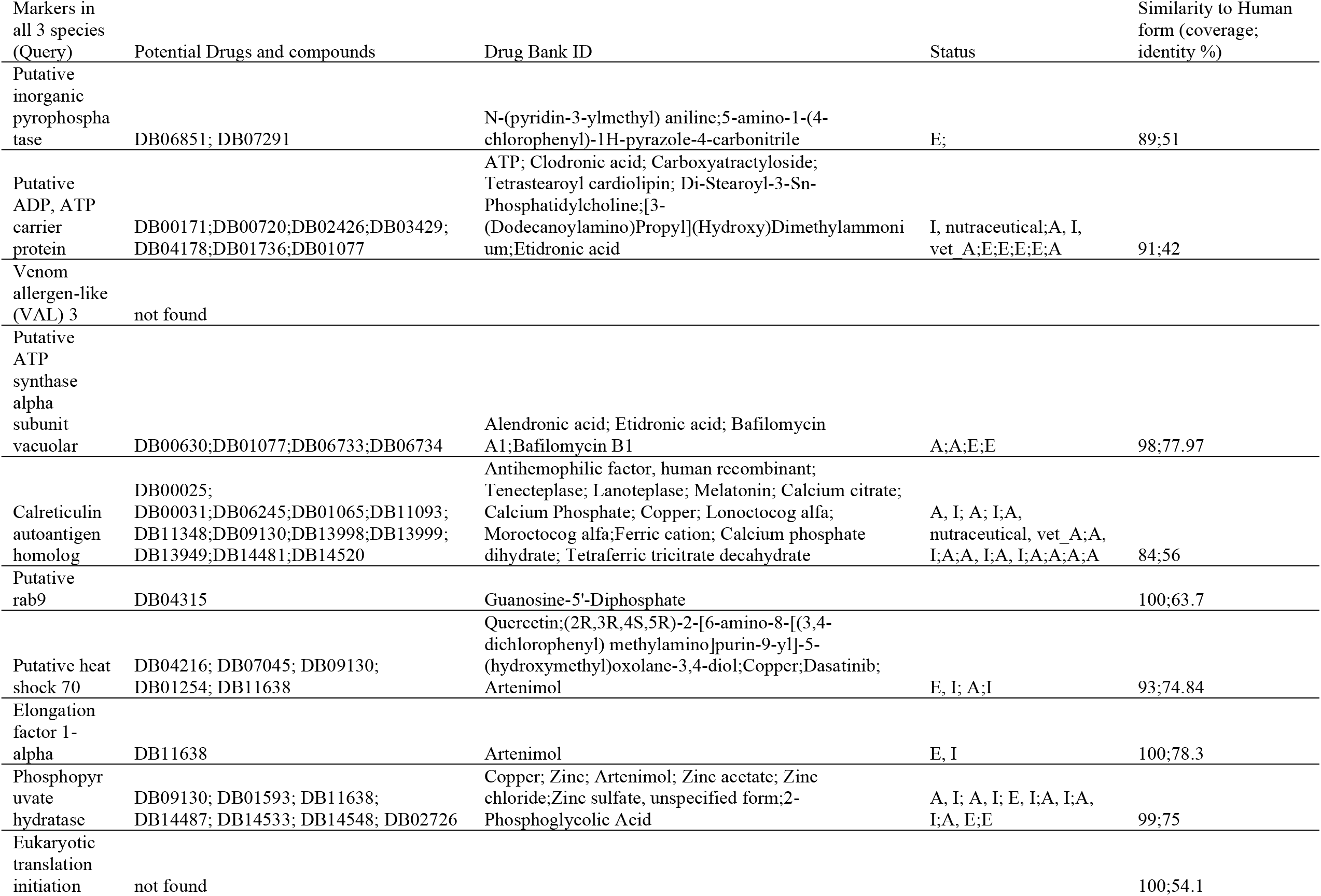
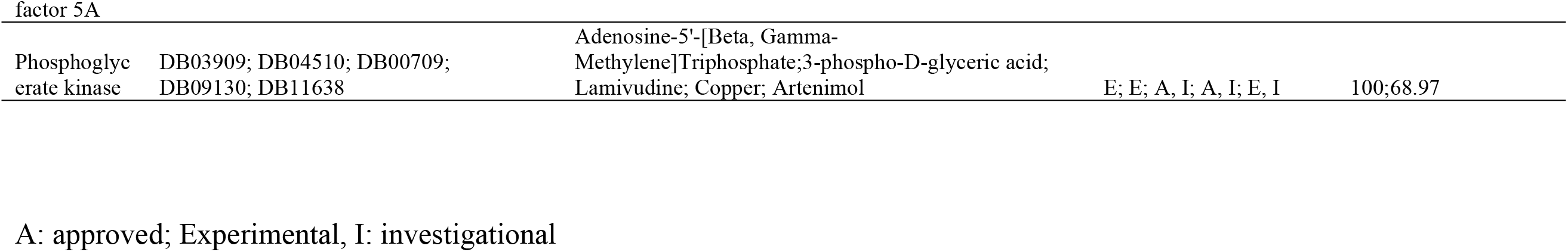
Drugs and compounds active against potential multi-species markers of medically important schistosomes

### Homology Determination

Of the 36 *S. mansoni* biomarker proteins (Table 2), 29 had identical proteins with both *Schistosoma haematobium* and *S. japonicum* or either of *S. haematobium* and *S. japonicum*. The biomarker proteins showed varying identities with *S. haematobium* and *S. japonicum* whole genome shotgun sequence while some of these proteins showed no significant similarity with either *S. haematobium* or *S. japonicum* whole genome shotgun sequence.

### Multiple Sequence Alignment

There were conserved regions in the Calreticulin autoantigen homolog (C4Q5I7) of *S. mansoni, S. haematobium* and of *S. japonicum* (AAC00515.1) showing that the amino acids in the protein sequences have strongly similar properties. The colors in the alignments show the locations of similarity and difference among the sequence based on the chemical nature of the amino acid residues (Figure 1).

**Figure 1.**
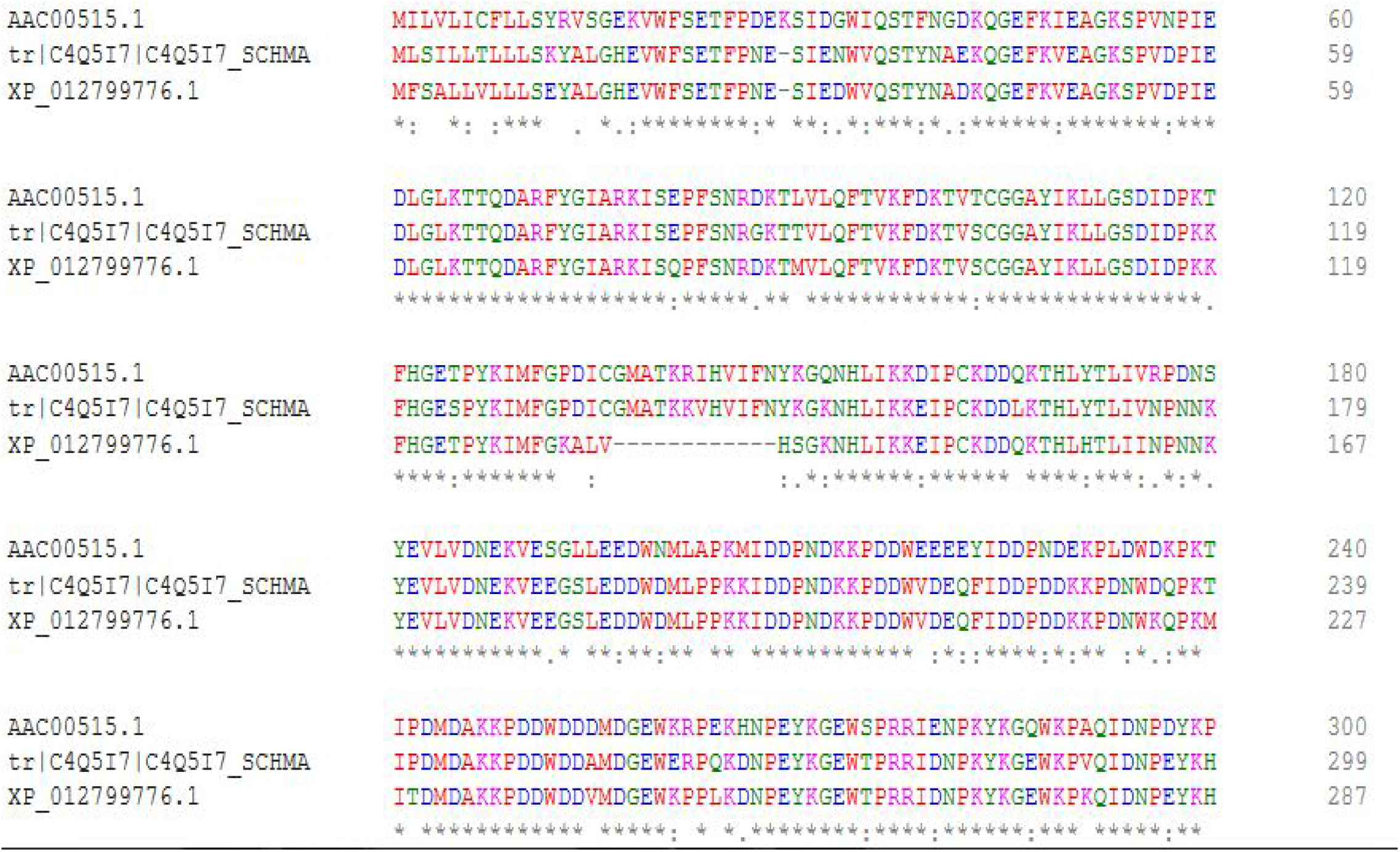
Calreticulin autoantigen homolog of *S. mansoni* (C4Q5I7); calreticulin of *S. haematobium* (XP_012799776.1) and calreticulin of *S. japonicum* (AAC00515.1) in multiple sequence alignment. “*” means that the amino acid in that column are identical in all sequence in the alignment. “:” denotes conserved regions i.e. amino acid with strongly similar properties. “.” denotes semi-conserved regions. The red color shows hydrophobic residues, blue color shows acidic residues, magenta shows basic residues, green shows hydroxyl residues while grey color show unusual amino acids. The colors in the alignments show the locations of similarity and difference among the sequence based on the chemical nature of the amino acid residues

Putative actin-1 of *S. mansoni* (G4VLW2) and the homologue Actin 5C (CAX75500.1) of *S. japonicum* shows conserved regions as well as regions of complete identical nucleotide sequence (Figure 2). Phosphoglycerate hydratase (enolase) showed conserved regions as well as regions of complete identical nucleotide sequence in most of the positions in the alignment (Figure 3)

**Figure 2.**
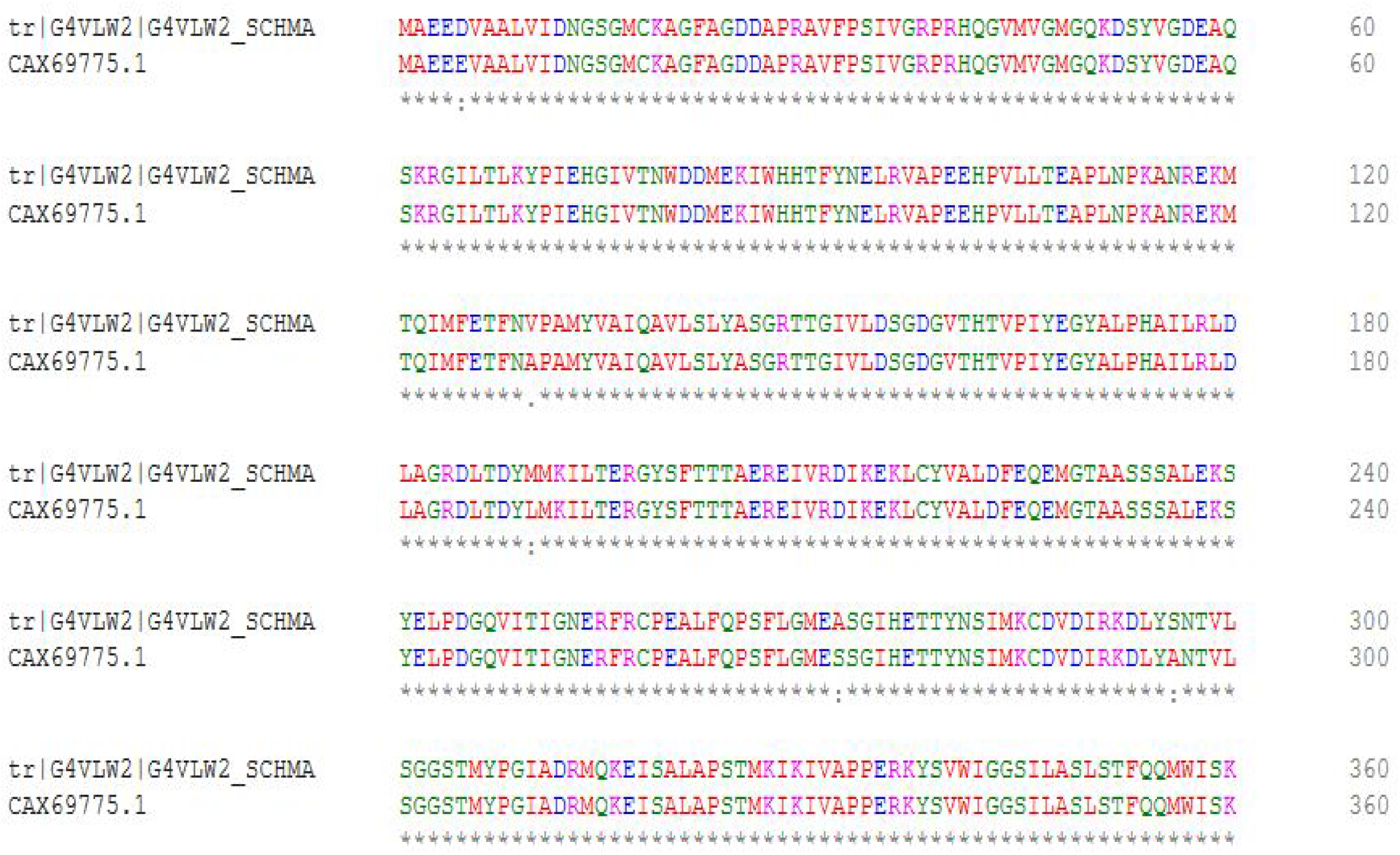
Multiple sequence alignment of putative actin-1 of *S. mansoni* (G4VLW2) and actin 5C of *S. japonicum* (CAX75500.1) “*” means that the amino acid in that column are identical in all sequence in the alignment. “:” denotes conserved regions i.e. amino acid with strongly similar properties. “.” denotes semi-conserved regions. The red color shows hydrophobic residues, blue color shows acidic residues, magenta shows basic residues, green shows hydroxyl residues while grey color show unusual amino acids. The colors in the alignments show the locations of similarity and difference among the sequence based on the chemical nature of the amino acid residues

**Figure 3.**
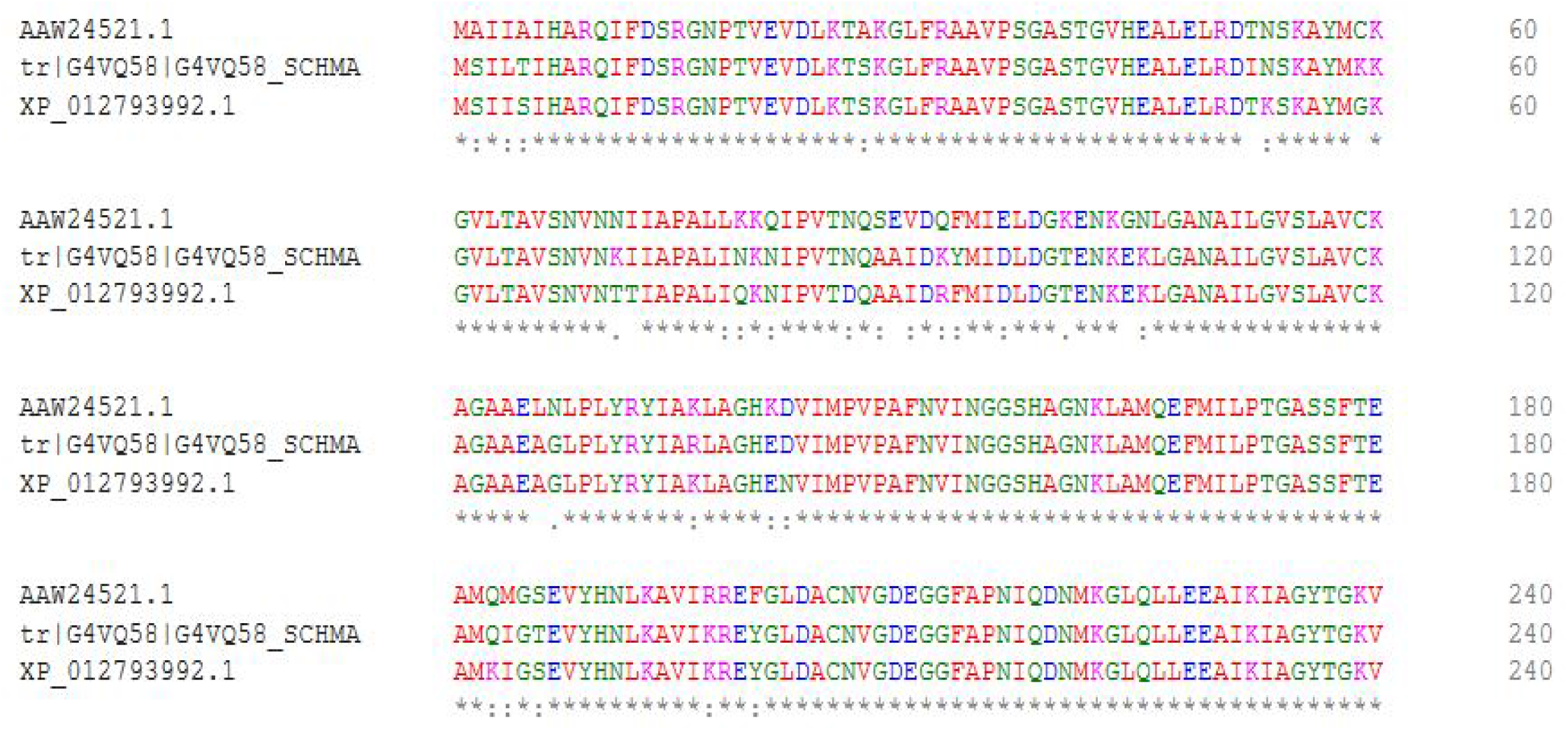
Multiple sequence alignment of phosphopyruvate hydratase of *S. mansoni* (G4VQ58); enolase of *S. haematobium* (XP_01293992.1) and SJCHGC00471 protein of *S. japonicum* (AAW24521.1) “*” denotes that the amino acid in that column are identical in all sequence in the alignment. “:” denotes conserved regions i.e. amino acid with strongly similar properties. “.” denotes semi-conserved regions. The red color shows hydrophobic residues, blue color shows acidic residues, magenta shows basic residues, green shows hydroxyl residues while grey color show unusual amino acids. The colors in the alignments show the locations of similarity and difference among the sequence based on the chemical nature of the amino acid residues

Most of the 29 *S. mansoni* biomarker proteins aligned with their homologues, giving rise to a shared identical conserved region. However, some vital biomarker proteins like venom allergen-like proteins which had been proposed as putative drug and vaccine target (Huang *et al*., 2017), showed more of semi conserved regions in which the amino acids had similar shape, but had weakly similar properties (Figure 4).

**Figure 4.**
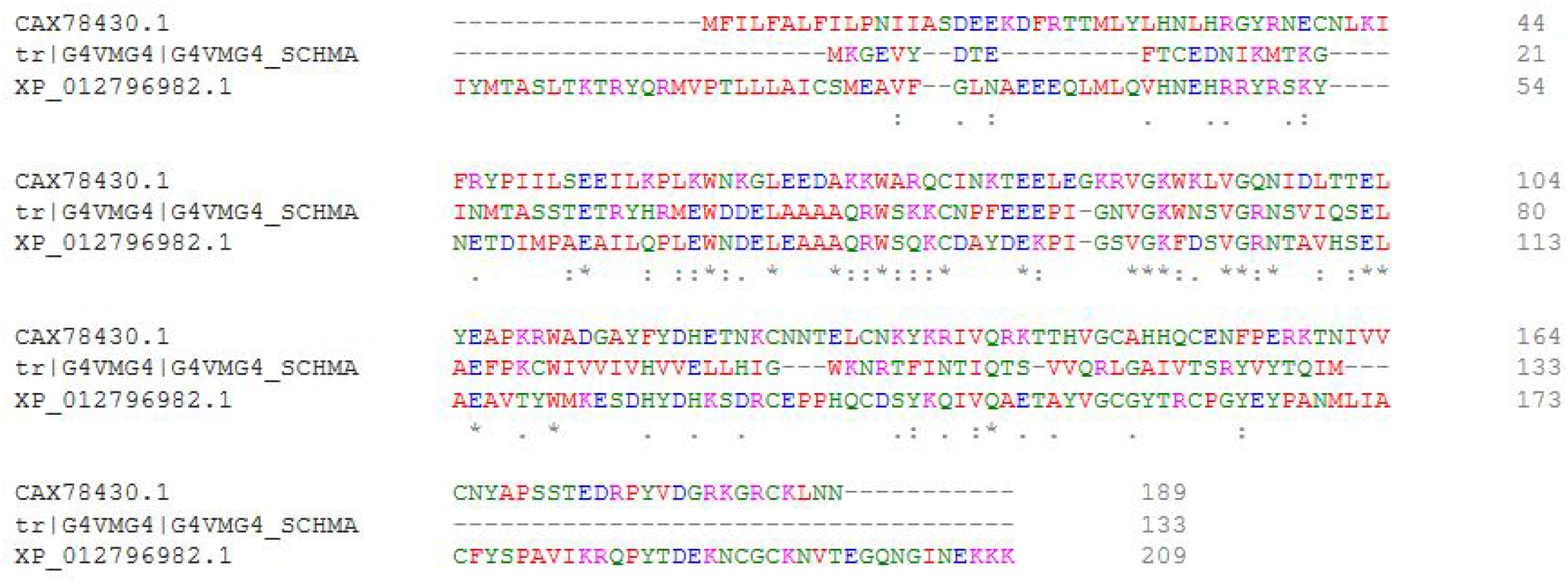
Multiple sequence alignment of venom allergen-like (VAL) 3 protein (G4VMG4) of *S. mansoni*; peptidase inhibitor 16, partial of *S. haematobium* (XP_012796982.1) and peptidase inhibitor 16,partial of *S. japonicum* (CAX78430.1) “*” means that the amino acid in that column are identical in all sequence in the alignment. “:” denotes conserved region i.e. amino acid with strongly similar properties. “.” denotes semi-conserved region. The red color shows hydrophobic residues, blue color shows acidic residues, magenta shows basic residues, green shows hydroxyl residues while grey color show unusual amino acids. The colors in the alignments show the locations of similarity and difference among the sequence based on the chemical nature of the amino acid residues

### Potential Therapeutic targets

For homologues identified consistently in all three species, 10 had potential targets known or being investigated in humans (Table 3). Such compounds included modified nucleotides, phosphatidylcholines, artenimol, and supplemental elements, among others.

## DISCUSSION

Schistosomiasis, a devastating and highly prevalent neglected tropical disease (NTD), is endemic in Nigeria (reviewed in Abdulkadir *et al*., 2017; Eke et al 2020). The emergence of schistosome parasites that are resistant to the traditional treatment due to over-reliance on praziquantel demands the development of newer and more effective treatment which requires accurate diagnostic methods (Fernandes *et al*., 2018). More needs to be known about the molecular pathology that influences the outcome of the infection by *S. haematobium* in order to develop new diagnostics, therapeutic and infection prevention strategies (Weerakoon *et al*., 2018).

The composition of the urine microbiome in schistosomiasis and its associated functions are being tackled by genomic sequencing technology. The identification of proteins common to all species of schistosomes will provide targets for developing drugs or vaccine to control schistosomiasis and the associated bladder cancer in view of the growing concern about resistance developing to the existing therapeutic (Magalhaes *et al*., 2016). The detection of cancer-associated biomarkers, preferably isolated from urine and blood, has therefore become important. Such biomarkers are now being developed and will provide tools that could be useful to evaluate the specific effects of long-term exposure to *S. haematobium*; cancer-specific urine biomarkers may therefore, play an important role in diagnosis of people with long-term *S. haematobium* infections (Sotillo *et al*., 2016).

This study was performed to establish homology of the 36 biomarker proteins evaluated in a previous study (Onile *et al*., 2017) in other schistosome species. It was found that protein homologues of the 36 *S. mansoni* biomarker proteins are present in *S. haematobium* and *S. japonicum*. Identification of homologues of these biomarker proteins could be used for early diagnosis of schistosomiasis and schistosomiasis associated bladder cancer and could also provide targets for developing drugs or vaccines that can be effective against all species of schistosomes.

The multiple sequence alignment was done by aligning *S. mansoni* biomarker proteins with their respective identical proteins retrieved. The basic information provided by the multiple sequence alignment is the identification of conserved sequence regions which is useful in designing wet laboratory experiment to test and modify the functions of these proteins, in predicting the function and structure of these proteins.

Calreticulin-like protein of *S. mansoni* which is a Ca2+ binding/storage protein, being found in a number of different animal taxa, and has been considered as a novel antigen for the detection of anti-*S. mansoni* antibodies (Wang *et al*., 2018). It has homologues in both *S. haematobium* and *S. japonicum* with many identical amino acids and conserved regions when aligned (Figure 1).

Putative actin -1 has been identified as a possible drug target for the treatment of schistosomiasis as a strong association between actin and *S. mansoni* adult worm surface membranes has been confirmed (Fernandes *et al*., 2011). Studies have described the role of actin in enhancing the activity of praziquantel (PQZ) treatment of schistosomiasis. It is suggested that PQZ intercalates in the surface membrane lipid bi-layers, thereby inducing tegumental changes that leads to antigen exposure, including actin. This study showed that putative actin-1 protein has a homolog in other *Schistosoma* species, with many identical regions and conserved regions (Figure 2).

Phosphopyruvate hydratase is homologous to enolase protein in *S. haematobium* (Figure 3), and is implicated in autoimmune diseases; its utilization by pathogens while invading host tissue is well documented. Therefore, it is a key player in understanding the host–parasite interaction, and it offers a target for chemo- and immunotherapy (Liu *et al*., 2014). Phosphopyruvate hydratase, elongation factor 1-alpha and putative actin-1 protein have been identified as eggshell protein markers that can induce cellular or antibody responses and may be useful schistosomiasis diagnostic candidates. They have also been identified as putative *S. mansoni* biomarker proteins (Onile *et al*., 2017). They all have homologues in both *S. haematobium* and *S. japonicum* with many conserved regions and semi conserved regions in the multiple sequence alignment (Figure 4.10).

HSP 60 is a molecular chaperon (Ludolf *et al*., 2014) which has a homologue in *S. haematobium*, though no significant similarity was found between the protein sequence and *S. haematobium* whole genome shotgun sequence; but no protein similar to HSP 60 was found in *S. japonicum* though it has significant similarity with *S. japonicum* whole genome shotgun sequence. The alignment of HSP 60 with its homologue in *S. haematobium* showed many conserved regions and regions where all the amino acids in the sequences were identical.

Venom allergen-like (VAL) proteins are associated with excretion/secretion products and extracellular environment of the parasite; they have been used as a trial vaccine against hookworm infections in humans (Wilbers et al, 2018). The VAL protein families are abundant in different helminth species including gastrointestinal nematodes, where they are known to carry out several roles in the infective activities of parasites (Tribolet *et al*., 2015). Venom allergen-like (VAL) 3 was identified among the proteins secreted in the tunnels formed by destruction of epidermal cells by cercariae, during the first two hours of penetration of human skin. Later, the protein was found to have the ability to bind lipids in vitro and complement the sterol export phenotype of yeasts in vivo (Fernandes et al., 2018). In this study, we showed that venom allergen-like protein has homologues in both *S. haematobium* and *S. japonicum* but with few conserved and semi-conserved regions (Figure 3).

Elongation factor 1-alpha, phosphopyruvate hydratase and histone-4 were all identified as potential *Schistosoma* biomarkers (Onile *et al*., 2017). These proteins have been identified previously in purified eggshell fragments of *Schistosoma mansoni* and also figured as schistosome antigens which may induce cellular or antibody responses (Dvorak *et al*., 2016). These eggshell markers may be very useful schistosomiasis diagnostic candidates rather than vaccine candidates, since such a vaccine would be likely to target the eggs and further encourage granuloma formation and pathology rather than priming the immune system against the parasite. These proteins all have homologues in other *Schistosoma* species with enormous conserved regions

Elongation factor 1-alpha has homologues in both *S. haematobium* and *S. japonicum* with many regions where the amino acids in the sequences are identical and many conserved regions occurring in the alignment of the proteins.

Lastly, of interest were the potential targets compounds for the homologues consistent in all three species. Although information from the drugbank database is often the result of testing/current testing on humans, some of the predictions could be of future use. For instance, Artenimol, which occurs often in Table 3, is being investigated for malaria treatment, as it is partly related to the Artemisinins which have been previously in small scale trials for parasitic disease treatment (https://clinicaltrials.gov/ct2/show/NCT02653898). A modified phosphatidylcholine was also a potential target for an ATP carrier protein (Table 3). This may make biological sense as we previously reported that changes in phosphatidylcholine metabolism was a hallmark of schistosomiasis (Adebayo et al, 2018). The nutraceutical elements are also interesting as they are cheaply available if they do have any benefits.

## CONCLUSION

Homology of the identified *S. mansoni* biomarker proteins was established in both *S. haematobium* and *S. japonicum*. This will help in early diagnosis of schistosomiasis and associated bladder cancer, although wet laboratory experiments are strongly indicated to serve as confirmation to the results of this study. This study can also help to provide targets for drugs or vaccines that can be simultaneously effective against all species of schistosomes.

## MATERIALS AND METHODS

### Data collection

Protein sequences of the 36 schistosome biomarkers were retrieved from the Uniprot database https://www.uniprot.org/ in FASTA format. *Schistosoma haematobium* and *S. japonicum* whole genome shotgun sequences were retrieved from GenBank https://www.ncbi.nlm.nih.gov/genbank/ as nucleotide and they were translated into protein format using EXPASY translating tool https://web.expasy.org/translate/.

## SMARTBLAST

The 36 *S. mansoni* biomarker proteins sequence in FASTA format were SMARTBLASTed https://blast.ncbi.nlm.nih.gov/smartblast/smartBlast.cgi to find protein sequences of other organism that are similar to the query protein sequence. The FASTA formats of protein sequences identical to each of the thirty-six protein were retrieved and used for multiple sequence alignment.

### Homology determination methods

The retrieved biomarker protein sequences were BLASTed against *Schistosoma haematobium* and *Schistosoma japonicum* translated whole genome shotgun sequence to search for homologous sequences. Sequence thus obtain were then aligned using BLASTP https://blast.ncbi.nlm.nih.gov to show the region of similarity.

### Multiple Sequence Alignment

Multiple alignments of protein sequences are important in studying sequences and providing basic information about conserved sequence region. Each of the thirty-six *S*.*mansoni* biomarker protein sequences were aligned with similar *S. haematobium* and *S. japonicum* protein sequences found when each of the protein was Smart Blasted. Multiple sequence alignment for each protein was done using Clustal Omega https://www.ebi.ac.uk/Tools/msa/clustalo/ which is a multiple sequence alignment program. The input protein sequences which were aligned were all in FASTA format.

### Potential Therapeutic targets

Markers with homologues in all three species were used as query on the drugbank database(drugbank.ca) in order to identify potential therapeutic targets which were already in use in humans or being investigated.

